# MEGAHIT k-mer range tuning trades computational efficiency for improved recovery of functional genes across cave sediment and wastewater metagenomes

**DOI:** 10.64898/2026.07.17.739120

**Authors:** Alina Cărunta, Horia Leonard Banciu, Alexandru Eugeniu Mizeranschi

## Abstract

Shotgun metagenomics is a powerful approach for profiling complex microbial ecosystems and discovering functional genes, including antimicrobial resistance genes (ARGs) and biosynthetic gene clusters (BGCs). De novo assembly with tools such as MEGAHIT commonly uses multiple k-mer lengths, but the effect of reduced k-mer sets on functional gene recovery has received limited attention. Here, we quantify the trade-off between assembly speed and functional-gene recovery using 17 cave sediment metagenomes and 10 wastewater metagenomes assembled under 19 MEGAHIT k-mer scenarios. In cave metagenomes, finer-grained k-mer ranges recovered more BGCs and, in several pairwise comparisons, more ARGs, but required longer runtimes. In wastewater metagenomes, finer-grained settings most clearly affected ARG recovery, whereas BGC counts did not differ significantly after Friedman testing. These results indicate that reduced k-mer sets can lower computational cost but may miss biologically relevant functional signal, depending on the dataset and downstream target. The study provides a quantitative basis for selecting MEGAHIT k-mer parameters according to whether computational efficiency or functional gene discovery is the primary aim.

## INTRODUCTION

Metagenomics (Oulas et al., 2015; Pérez-Cobas et al., 2020) is a field of study that allows researchers to explore the vast microbial diversity present in diverse environments, including surface and subsurface soil and water habitats, man-made environments or human-associated microbiomes (Guo et al., 2015; Che et al., 2019; Turrini et al., 2020; Chiciudean et al., 2022; Vojvoda Zeljko et al., 2024; Gorecki et al., 2024). By analyzing DNA sequences directly extracted from these complex samples, metagenomics enables the identification and characterization of novel microbial species and their functional potential. Metagenomic assembly is a powerful method for exploring whole microbial communities, including species which cannot be cultivated in a laboratory. Specialized data analysis tools and pipelines enable the prospecting of microbial species for possibly novel functional genes and study their encoded products and associated metabolic traits. Such novel genes of interest include antimicrobial resistance genes (ARGs) (Tenover, 2006) and biosynthetic gene clusters (BGCs) (Cimermancic et al., 2014), with a broad impact on understanding microbial interactions, dispersal and occurrence of antimicrobial resistance, or prospective biotechnological applications.

ARGs confer resistance to various antimicrobials upon their host microorganisms (Aminov and Mackie, 2007). BGCs, also known as metabolic gene clusters, consist of sets of non-homologous genes, which are involved in metabolic pathways (Lebedeva et al., 2021). BGCs are predicted to play key roles in defense mechanisms against antimicrobials. Another role of BGCs is the production of secondary metabolites involved in microbial competition (Rutledge and Challis, 2015). Consequently, the screening of ARGs and BGCs across diverse microbial communities is considered a promising approach to facilitate discoveries with potential implications for innovative industrial bioprocesses (Zada et al., 2022).

In recent years, several pipelines were developed for metagenomic assembly, binning and functional annotation using domain-specific languages such as Nextflow (Di Tommaso et al., 2017), which was used to build the nf-core family of data analysis pipelines (Ewels et al., 2020). In the field of metagenomics, relevant examples of pipelines include *nf-core/mag* (Krakau et al., 2022) for assembly and binning, and nf-core/funcscan (Yates et al., 2024) for functional annotation, including gene annotation and the discovery of ARGs and BGCs. These pipelines automate the specialized workflows from raw sequenced reads to assembled and annotated (meta)genomes. Once configured, they reproducibly generate assembly and annotation outputs from any input data, greatly simplifying re-analysis.

A crucial step in any metagenomics pipeline is the metagenome assembly of raw sequenced reads into contiguous sequences or contigs. Numerous tools and approaches have been proposed for this and several studies evaluated the performance of these tools (Van der Walt et al., 2017; Vollmers et al., 2017; Sczyrba et al., 2017; Ayling et al., 2020; Yang et al., 2021). Software packages such as metaSPAdes (Nurk et al., 2017) and MEGAHIT (Li et al., 2015) are usually recommended, the former generally having better accuracy, while the latter offers better efficiency (considerably smaller runtimes and memory requirements), without bringing a significant drop in accuracy.

For example, Ayling et al. (2020) proposed new approaches for metagenome assembly with short reads, by describing the existing approaches for metagenome assemblers, including tools like MEGAHIT and metaSPAdes. Van der Walt et al. (2017) compared the performance of various metagenome assemblers on complex metagenomes. The authors provided a workflow that aims to guide researchers on selecting the best metagenome assembler for their metagenomic project, taking into account several aspects, including available computational resources. Among their recommendations, the MEGAHIT software package was highlighted due to several advantages over competing tools. Similar to Van der Walt et al. (2017), Vollmers et al. (2017) compared and evaluated the performance of several metagenome assembly tools, including MEGAHIT, on metagenomic samples, from the perspective of a microbiologist. Sczyrba et al. (2017) performed a critical evaluation of the metagenome interpretation for several metagenomic software tools, including MEGAHIT. Finally, Yang et al. (2021) surveyed several computational tools, including MEGAHIT, used to generate metagenome-assembled genomes (MAGs) from metagenomic sequencing data, and provided guidelines for selecting the appropriate tool for MAG generation.

Several advantages of MEGAHIT were repeatedly highlighted in the previously mentioned studies: it requires less time and less memory compared to SPAdes for assembling complex datasets; it is an open-source tool; it offers a relatively high flexibility; it provides the highest assembly cost-efficiency trade-off; it has high sensitivity, i.e., a generally higher probability to include low-abundant community members in the assemblies; and it is very well documented.

A common strength of both metaSPAdes and MEGAHIT is that they rely on a multiple k-mer-based approach. Assemblers that use multiple k-mer sizes generally have a better performance compared to those assemblers using a single k-mer size (Ayling et al., 2020). For example, setting values for the k-mer size range in the MEGAHIT assembler is possible via parameters *k-min, k-max* and *k-step*, where *k-min* is the minimum k-mer size, *k-max* is the maximum k-mer length and *k-step* is an incremental step. These 3 parameters can effectively specify a range of k-mer sizes, for which MEGAHIT creates a set of succinct de Bruijn graphs which are then integrated to form the final assembly. For example, Vollmers et al. (2017) have used a k-mer range defined by *k-min*=21, *k-max*=101 and *k-step* of 10, which yields the following set of k-mer sizes: 21, 31, 41, 51, 61, 71, 81, 91 and 101. If the desired k-mer range cannot be represented by such an arithmetic progression, it can be explicitly defined using the *k-list* parameter.

In the context of metagenome assemblers that employ multiple k-mer sizes, the relationship between the k-mer sizes and the quality of resulting assemblies has generally been unexplored. An exception is the work of Qayyum et al. (2025), who compared three scenarios related to k-mer sizes in MEGAHIT and found that a reduced set of k-mer sizes produced similar quality metrics as those from larger (more fine-grained) sets of k-mers, while bringing about a considerable reduction in assembly time. However, in this study, the impact of the k-mer size settings on the number of identified ARGs and BGCs was unexplored.

Another previous study from Cărunta et al. (2024) tackled the three previously mentioned assembly parameters on several quality metrics, including the number of ARGs and BGCs, using multiple assembly scenarios (based on different k-mer size ranges) and 7 samples extracted from Movile Cave in Romania. Their findings refuted those of Qayyum et al. (2025) regarding the use of the reduced set of k-mer sizes when the target is the identification of ARGs and BGCs. According to the results of Cărunta et al. (2024), the use of the reduced set of k-mer sizes led to the discovery of fewer ARGs and BGCs compared to the case when using a larger set of k-mer sizes.

The objective of the current study is to extend the results of Cărunta et al. by targeting the limitations identified within the previously mentioned study (Cărunta et al., 2024). Specifically, we include additional sediment samples extracted from three new cave sites, and expand the set of assembly scenarios to be compared, including new scenarios with low and high values for the *k-step* parameter. In this study, we devised a comprehensive set of scenarios related to the three parameters *k-min, k-max* and *k-step* and compared the resulting assemblies on various metrics related to assembly quality, as well as the number of identified ARGs and BGCs and their lengths expressed in base pair counts. We also focused on comparing only the pairs of scenarios in which the value of a single assembly parameter (*k-min, k-max* or *k-step*) varied, making it easier to interpret their effects. In addition to the cave samples, a second dataset for wastewater metagenomes (Fierer et al., 2022) was also used to test the generalisability of our results on an independent dataset.

## METHODS

The data on which this study was conceived originated from 17 cave samples as follows: 5 sediment samples from Cloşani Cave, 4 sediment samples from Ferice Cave, one sediment sample from Bears’ Cave, and 7 sediment samples collected from Movile Cave in Romania, as previously described (Chiciudean et al., 2022). These samples were sequenced and paired-end reads were generated using PE150 on the Illumina NovaSeq 6000 platform. The raw sequenced reads that were generated from these samples were uploaded to the NCBI SRA database under the project accession number PRJNA777757. The per-sample accession IDs are listed in Table S1. A second dataset for wastewater metagenomes from a wastewater system on a university campus was also included (Fierer et al., 2022). Specifically, this dataset was originally published under NCBI project accession number PRJNA875025 and contains 188 wastewater samples collected from 17 locations on the University of Colorado campus. A subset of 10 wastewater samples was randomly selected for this study. The individual accession IDs, sample IDs, read-pair counts and base counts for these 10 samples are provided in Table S2.

The *nf-core/mag* pipeline v. 3.0.2 (Krakau et al., 2022) was used for metagenome assembly and binning. Pipeline executions were set up for each of the assembly scenarios depicted in Table 1. These assembly scenarios are represented as tuples of values for each of the three k-mer parameters (i.e., *k-min, k-max, k-step*), taking into consideration the general constraints imposed by MEGAHIT for these parameters, as follows. The parameter *k-min* must be an odd integer ≥15. The parameter *k-max* must be an odd integer ≤127, but greater than or equal to *k-min*. The parameter *k-step* must be an even integer ≥2 and ≤28, such that all k-mer sizes are odd, within a valid range from 15 to 127. MEGAHIT v. 1.2.9 (Li et al., 2015) was used for the metagenome assembly and quality metrics were collected from its logs. In order to increase specificity, mercy k-mers were excluded and only contigs of a minimum length of 2000 bp were retained. The values for the k-mer length parameters (*k-min, k-max* and *k-step*) were configured as depicted in Table 1.

**Table 1.** Scenarios for assembly generation in MEGAHIT.

| scenario | k-min | k-max | k-step |
| --- | --- | --- | --- |
| k21_121_04 | 21 | 121 | 04 |
| k21_121_10 | 21 | 121 | 10 |
| k21_121_20 | 21 | 121 | 20 |
| k21_111_10 | 21 | 111 | 10 |
| k21_101_04 | 21 | 101 | 04 |
| k21_101_10 | 21 | 101 | 10 |
| k21_101_20 | 21 | 101 | 20 |
| k31_121_10 | 31 | 121 | 10 |
| k31_111_04 | 31 | 111 | 04 |
| k31_111_10 | 31 | 111 | 10 |
| k31_111_20 | 31 | 111 | 20 |
| k31_101_10 | 31 | 101 | 10 |
| k41_121_04 | 41 | 121 | 04 |
| k41_121_10 | 41 | 121 | 10 |
| k41_121_20 | 41 | 121 | 20 |
| k41_111_10 | 41 | 111 | 10 |
| k41_101_04 | 41 | 101 | 04 |
| k41_101_10 | 41 | 101 | 10 |
| k41_101_20 | 41 | 101 | 20 |

In contrast to the previous study by Cărunta et al. (2024), we excluded the comparison of the resulting assemblies on metrics related to the number of metagenomic bins (e.g., *filt bins, unfilt bins*). These quality metrics related to metagenomic bins were excluded here because they did not have any relationship with the identification of ARGs and BGCs, these functional genes being searched within all contigs of samples and thus were not affected by the binning process. Another reason for excluding these metrics is the fact that in the previous study they did not have an overall significant effect on the three assembly parameters (Cărunta et al., 2024). This exclusion of quality metrics thus reduced the complexity of the analyses performed within this study. Note that the terms “quality metric” and “variable of interest” are used interchangeably within this study.

The sample-specific assemblies created with MEGAHIT were given as input to *nf-core/funcscan* (Yates et al., 2024). The version v. 2.1.0 of the pipeline was used for this study. A single analysis encompassing the assemblies created from all the scenarios was set up, enabling the identification of ARGs using AMRFinderPlus v. 3.12.8 (Feldgarden et al., 2021) and the identification of BGCs using GECCO v. 0.9.10 (Carroll et al., 2021).

Spearman’s rank correlations between various variables (from Table 2) were calculated to study the strength and type of their association (positive vs. negative correlation). Furthermore, Friedman’s test followed by Conover’s all-pairs comparisons test for unreplicated blocked data were applied to identify significant differences between the different scenarios, considering 11 dependent variables that were collected within this study (Table 2). Statistical analyses were performed in the R programming language v. 4.4.3 (R Core Team, 2024) using the friedman.test function from the stats package and the frdAllPairsConoverTest function from the PMCMRplus package v. 1.9.12 (Pohlert, 2023), with Bonferroni correction of p-values for multiple comparisons. The results were considered statistically significant at a p-value threshold of 0.05. Hierarchical clustering using Euclidean distance was performed on Conover’s test statistics with the hclust function from the stats package with the ward.D2 method, in order to underline similarity patterns within the scenario pairs and variables, respectively.

**Table 2.** Definitions for quality metrics used in this study. Up arrow symbol (*↗*) means larger values are better than smaller values for the corresponding quality metric, while using the down arrow symbol (*↘*) means the opposite.

| Metric | Definition | Target |
| --- | --- | --- |
| megahit_s | the assembly runtime in MEGAHIT, expressed in seconds | ↘ |
| contigs | the number of contigs in the corresponding assembly | ↗ |
| rRNA_genes | the number of identified rRNA genes | ↗ |
| L50 | the minimum number of contigs, ordered by decreasing length, needed to cover 50% of the total assembly length | ↘ |
| max_contig | the length of the largest contig | ↗ |
| N50 | the contig length for which contigs of that length or longer account for at least 50% of the total assembly length | ↗ |
| total_bp | the total assembly size in bp | ↗ |
| ARG | the number of identified ARGs | ↗ |
| ARG_len | the mean of gene lengths per sample per scenario | ↗ |
| BGC | the number of identified BGCs | ↗ |
| BGC_len | the mean of the BGC lengths per sample per scenario | ↗ |

All analyses were performed on a virtual computing cluster built on top of OpenStack using HTCondor v. 8.8.15 (Bockelman et al., 2021), with NFS storage and 8 execution nodes, each with 64 AMD EPYC 7702 2.0 GHz vCPUs and 256 GB of memory. In order to ensure an adequate comparison of runtimes across the different scenarios, dedicated access to the cluster was ensured and each MEGAHIT process was allowed to use the full amount of vCPUs and memory available on a single cluster node.

## RESULTS

A total of 19 assembly scenarios were analyzed in this study and the corresponding values for the parameters *k-min, k-max* and *k-step* in MEGAHIT are presented in Table 1. The different settings for these scenarios were chosen as variations from the one employed by Vollmers et al. (2017) (i.e., *k-min*=21, *k-max*=101 and *k-step*=10). These scenarios were tested on both previously mentioned datasets. From the scenario-specific results of the *nf-core/mag* and *nf-core/funcscan* pipelines, the following quality metrics were collected and analyzed: *megahit s, contigs, rRNA genes, L50, max contig, N50, total bp, ARG, ARG len, BGC* and *BGC len*. Specifically, information related to the first seven variables (*megahit s, contigs, rRNA genes, L50, max contig, N50* and *total bp*) was provided by MEGAHIT. For the last four variables, namely *ARG, ARG len, BGC* and *BGC len*, data were collected from the results of the *nf-core/funcscan* pipeline. These metrics are defined in Table 2.

## Results on cave metagenome samples

Figure 1 depicts Spearman’s rank correlations between the different variables of interest. While roughly half of the correlations were mild (with values between −0.4 and 0.4), several pairs of variables with a strong positive correlation (values above 0.9) were encountered in Figure 1. For example, the assembly length (*total bp*) and the contig count (*contigs*) were highly correlated (*r*_*S*_ ≈ 0.98) and both were strongly correlated with the BGC count (*r*_*S*_ ≈ 0.90 and *r*_*S*_ ≈ 0.82, respectively), indicating that larger assemblies typically yielded more predicted BGCs (Figure 1).

**Figure 1.**
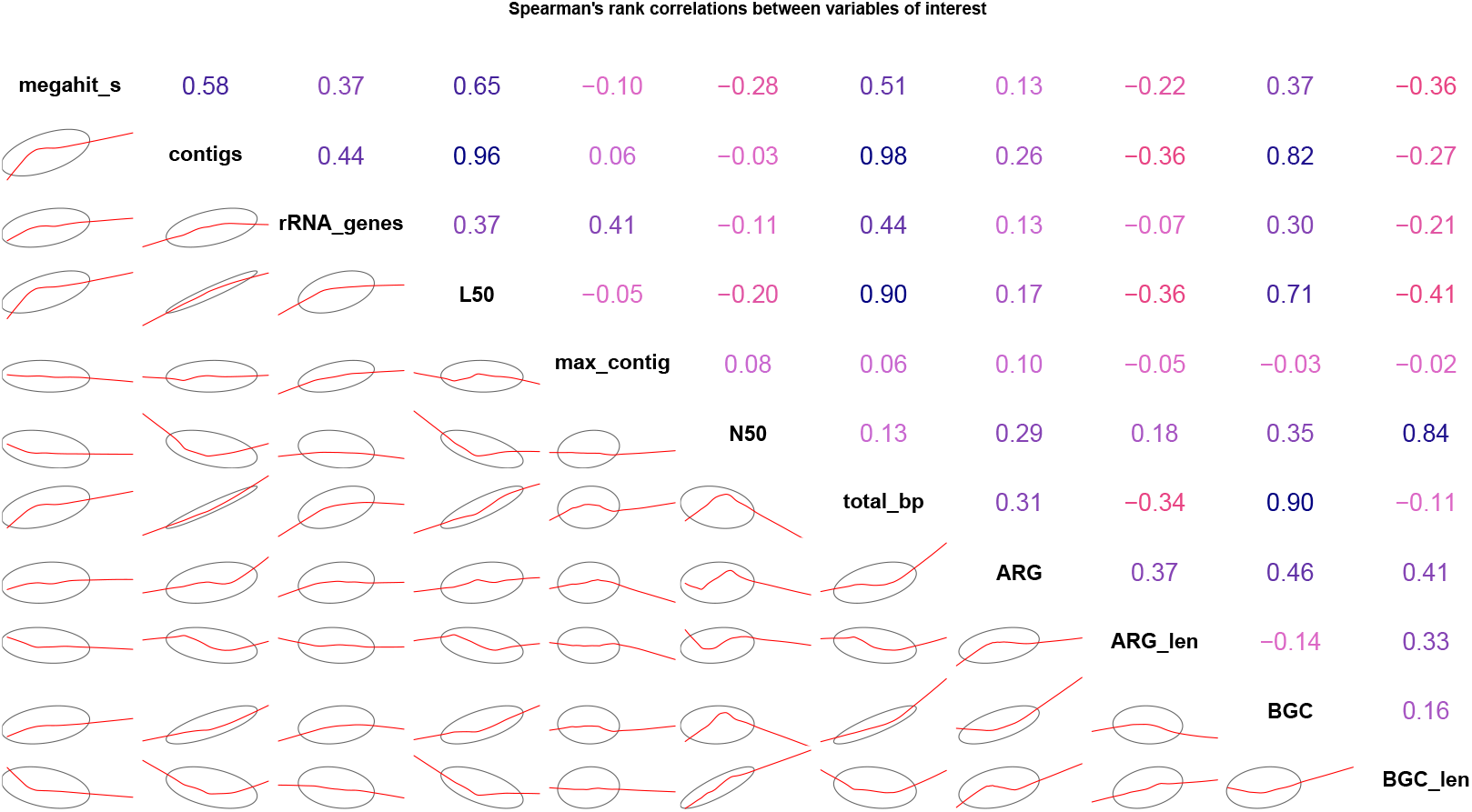
Spearman’s rank correlations between variables using the cave dataset. Variables are on the main diagonal; variable correlation coefficients for the whole dataset are on the upper-right panel; confidence ellipses and smoothed red lines, based on the corresponding correlation coefficient, indicating the overall 2D shape for each pair of two variables are shown on the lower-left panel. Blue color indicates positive correlation and red color indicates negative correlation, whereas the color intensity signifies the strength of the correlation.

We performed statistical analyses using Friedman’s test based on the results of the two previously-mentioned pipelines (i.e., *nf-core/mag* and *nf-core/funcscan*) (see Table 3), in order to identify statistically significant differences for the eleven variables across the different pairs of scenarios. We applied Friedman’s test using samples as blocks and scenarios as groups. The variable *ARG len* was removed from further analysis because its corresponding p-value did not pass the significance threshold in Friedman’s test.

**Table 3.** P-values for assembly quality metrics computed using Friedman’s test for the cave dataset. The values were rounded to 3 decimals.

| Variable | p_value_Friedman |
| --- | --- |
| megahit_s | <0.001 |
| contigs | <0.001 |
| rRNA_genes | <0.001 |
| L50 | <0.001 |
| max_contig | <0.001 |
| N50 | <0.001 |
| total_bp | <0.001 |
| ARG | <0.001 |
| ARG_len | 0.487 |
| BGC | <0.001 |
| BGC_len | <0.001 |

Next, we applied Conover’s all-pairs comparisons test for unreplicated blocked data on the remaining ten variables (Table S7). Similar to Friedman’s test, we set up samples as blocks and scenarios as groups, with the Bonferroni correction being used for the p-value adjustment. The tests generated p-values for 171 possible pairwise scenario comparisons, but only 39 such comparisons were considered for further analysis, where only one of the three k-mer length parameters was allowed to vary between the compared scenarios, while the other 2 parameters were kept constant. The selected 39 comparative scenarios are listed in Tables S3, S4 and S5. The rest of 132 pairwise scenario comparisons (where two or more k-mer length parameters were simultaneously distinct between the two compared scenarios) were discarded because they were difficult to interpret in the study’s context, focusing on specific k-mer parameter effects.

The results of Conover’s all-pairs comparisons test for unreplicated blocked data are summarized in Fig. 2 in the form of a heatmap. The corresponding values of Conover’s significant statistics for all ten variables can be found in Table S7. Of the ten variables that were analyzed, *max_contig* showed significant differences only in six of the total of thirty-nine pairs of scenarios. Similarly, *BGC_len* showed significant pairwise differences in five of the thirty-nine scenario pairs, while *BGC* showed significant pairwise differences in thirty-two of the thirty-nine scenario pairs. The *contigs* and *total_bp* variables had significantly different rank sum values in thirty-three and thirty-four of the thirty-nine scenario pairs, respectively. The variable *megahit_s* was significantly different in all but five of the thirty-nine scenario pairs, just as the variable *total_bp*. Furthermore, the other variables (*rRNA_genes, N50, L50* and *ARG*) showed significant differences in various numbers of scenario pairs, namely: twenty-two, twenty, nineteen and fourteen pairs, respectively. Thus, these gene-related metrics showed more moderate changes across k-mer settings than assembly size or runtime.

**Figure 2.**
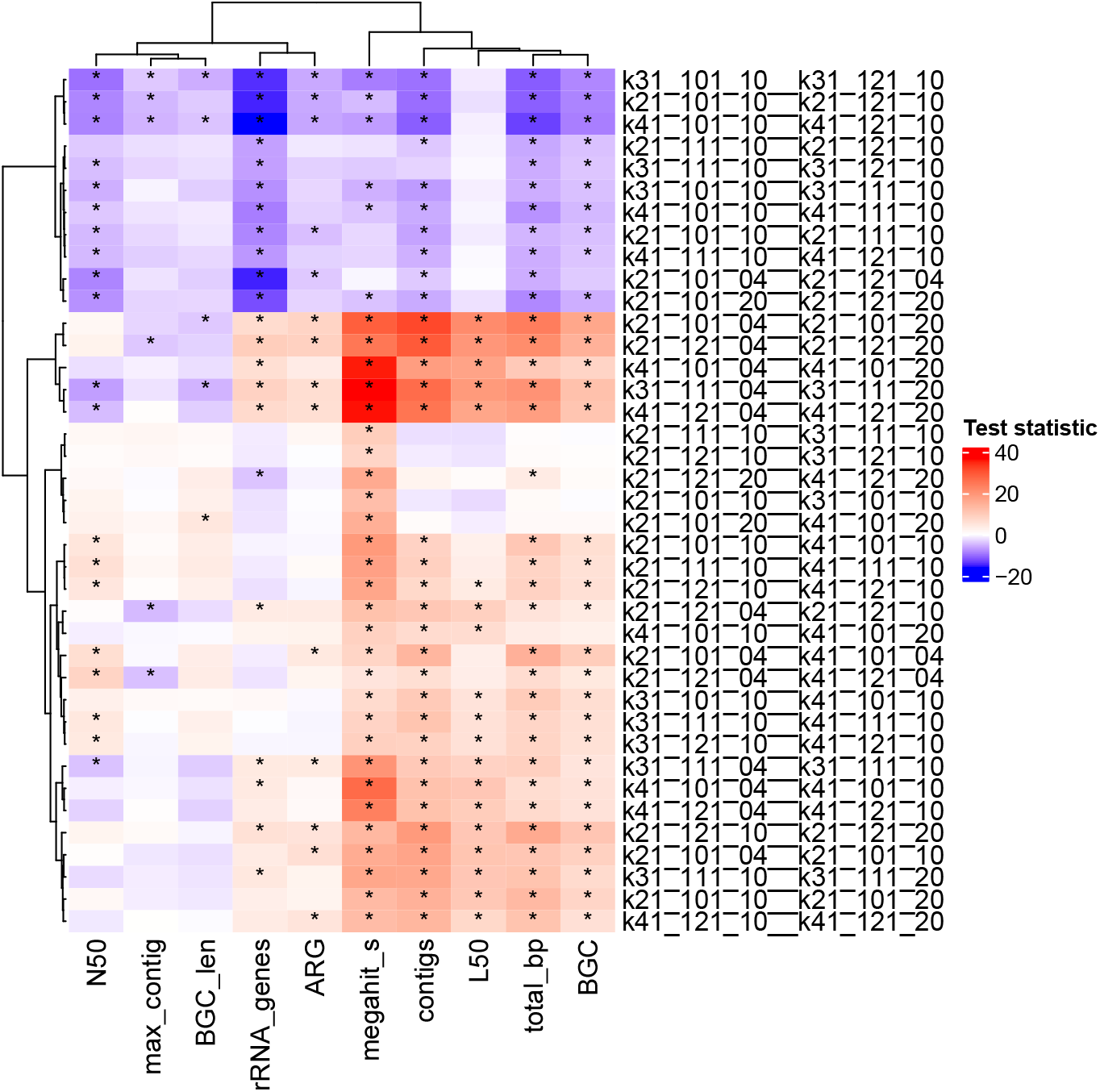
Heatmap showing test statistics from Conover’s all-pairs comparisons test for unreplicated blocked data, assessing differences across ten variables and thirty-nine pairwise scenario comparisons for the cave dataset. For example, scenario pair k21_101_04_k21_101_10 can be translated as a comparison between scenario k21_101_04 vs. scenario k21_101_10 when only parameter k-step varies. Red shades indicate that the first scenario in a pair had a higher rank sum (positive test statistic), while blue shades indicate the opposite (negative test statistic). Comparisons with p-values ≤ 0.05 are marked with asterisks. Rows (scenario pairs) and columns (variables) are hierarchically clustered using Euclidean distance to highlight patterns in statistical differences.

Regarding the hierarchical clustering of the pairwise scenario comparisons depicted on rows in Fig. 2, four main clusters were noticeable. These clusters of scenario pairs are presented in Table 4, where each column contains a list of scenario pairs that belong to a certain cluster. Specifically, cluster 1 contains pairwise scenarios with varying *k-max*, Cluster 2 consists of pairwise scenarios with different *k-step* values, Cluster 3 is composed of pairwise scenarios with different *k-min*, while cluster 4 consists of pairs where either *k-min* or *k-step* varied.

**Table 4.** List of scenario pairs clustered using Euclidean distance for the cave dataset. Each column contains scenario pairs that belong to a specific cluster and only one k-mer parameter varies within each pair.

| Cluster 1<br>(k-max) | Cluster 2<br>(k-step) | Cluster 3<br>(k-min) | Cluster 4<br>(k-min \| k-step) |
| --- | --- | --- | --- |
| k31_101_10_k31_121_10 | k21_101_04_k21_101_20 | k21_111_10_k31_111_10 | k21_101_10_k41_101_10 |
| k21_101_10_k21_121_10 | k21_121_04_k21_121_20 | k21_121_10_k31_121_10 | k21_111_10_k41_111_10 |
| k41_101_10_k41_121_10 | k41_101_04_k41_101_20 | k21_121_20_k41_121_20 | k21_121_10_k41_121_10 |
| k21_111_10_k21_121_10 | k31_111_04_k31_111_20 | k21_101_10_k31_101_10 | k21_121_04_k21_121_10 |
| k31_111_10_k31_121_10 | k41_121_04_k41_121_20 | k21_101_20_k41_101_20 | k41_101_10_k41_101_20 |
| k31_101_10_k31_111_10 |  |  | k21_101_04_k41_101_04 |
| k41_101_10_k41_111_10 |  |  | k21_121_04_k41_121_04 |
| k21_101_10_k21_111_10 |  |  | k31_101_10_k41_101_10 |
| k41_111_10_k41_121_10 |  |  | k31_111_10_k41_111_10 |
| k21_101_04_k21_121_04 |  |  | k31_121_10_k41_121_10 |
| k21_101_20_k21_121_20 |  |  | k31_111_04_k31_111_10 |
|  |  |  | k41_101_04_k41_101_10 |
|  |  |  | k41_121_04_k41_121_10 |
|  |  |  | k21_121_10_k21_121_20 |
|  |  |  | k21_101_04_k21_101_10 |
|  |  |  | k31_111_10_k31_111_20 |
|  |  |  | k21_101_10_k21_101_20 |
|  |  |  | k41_121_10_k41_121_20 |

A similar situation to pairwise scenario clustering was encountered with the clustering of columns containing variables of interest, where two main clusters were detected. The first cluster was composed of variables *N50, max_contig, BGC_len, rRNA_genes* and *ARG*. Meanwhile, the second cluster consisted of the variables *megahit_s, contigs, L50, total_bp* and *BGC*.

In order to ease the identification of significant differences in Fig. 2, the p-values for pairwise comparisons that were greater than 0.05 have been omitted from the figure. The p-values for pairwise comparisons that were less than or equal to 0.05 are marked with asterisks.

The impact of different settings for the *k-step* parameter (values 4, 10, 20) on several quality metrics is presented in Fig. 3. Within this figure, each panel shows the *k-step* trade-off regarding a specific variable of interest, namely (a) N50 lengths, (b) assembly runtimes in seconds for MEGAHIT, (c) BGC counts and (d) ARG counts, based on their mean value grouped by *k-step*. Similarly, the impact of different settings for the *k-min* parameter (values 21, 31, 41) on the same quality metrics was shown in Fig. 4. Further, the impact of different settings for the *k-max* parameter (values 101, 111, 121) on each of the four quality metrics (i.e., *N50, megahit_s, BGC* and *ARG*) is presented in Fig. 5.

**Figure 3.**
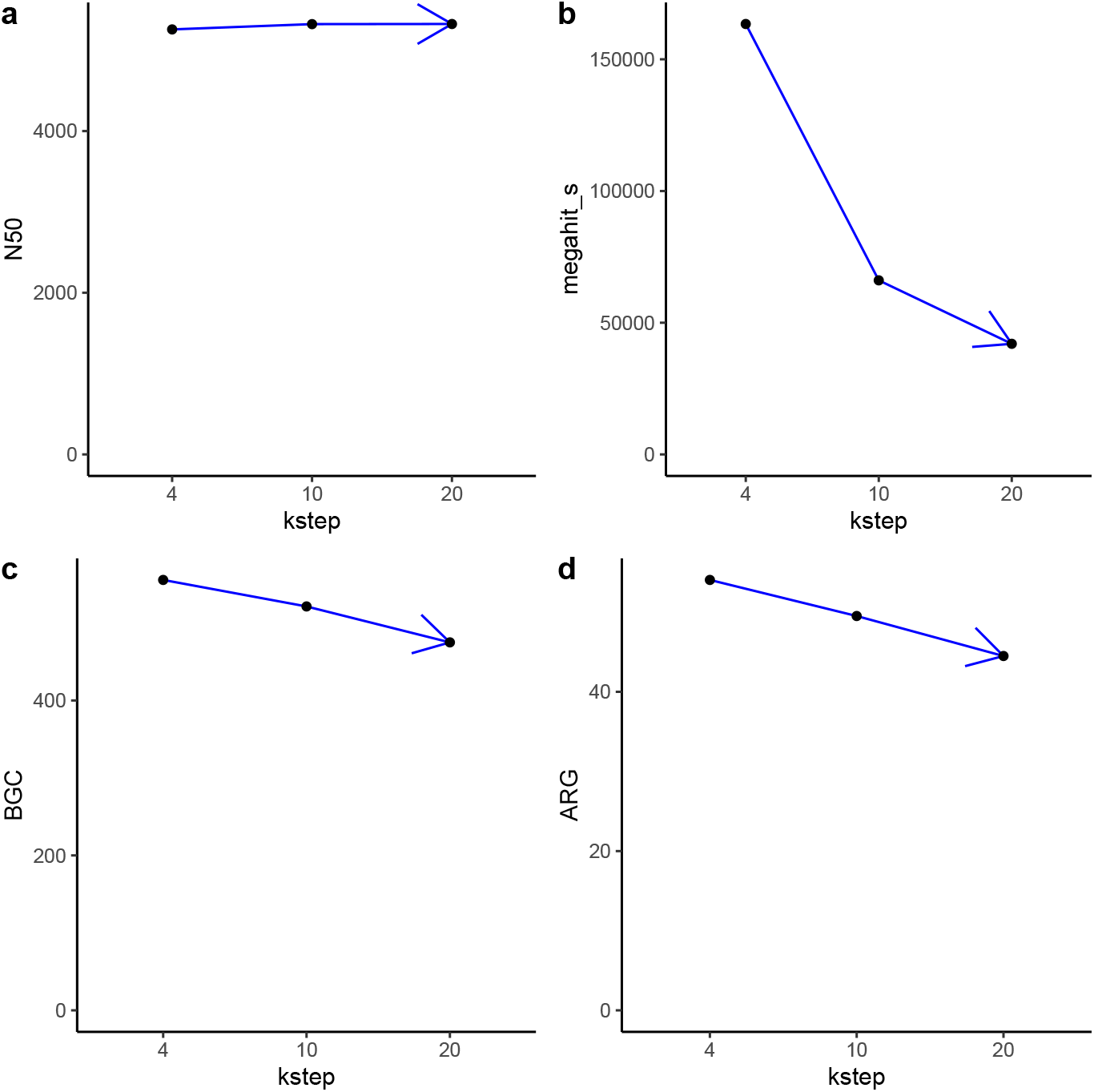
The *k-step* trade-off using the cave dataset where the x-axis contains values for the *k-step* parameter (i.e., 4, 10 and 20). Values from the y-axis from each panel correspond to the mean of a particular variable of interest grouped by the values of the *k-step* parameter. **a** Comparison between the mean of N50 lengths by different *k-step* settings (left-top corner). **b** Comparison between the mean of assembly runtimes in seconds using MEGAHIT by different *k-step* settings (right-top corner). **c** Comparison between the mean of BGC counts by different *k-step* settings (left-bottom corner). **d** Comparison between the mean of ARG counts by different *k-step* settings (right-bottom corner).

**Figure 4.**
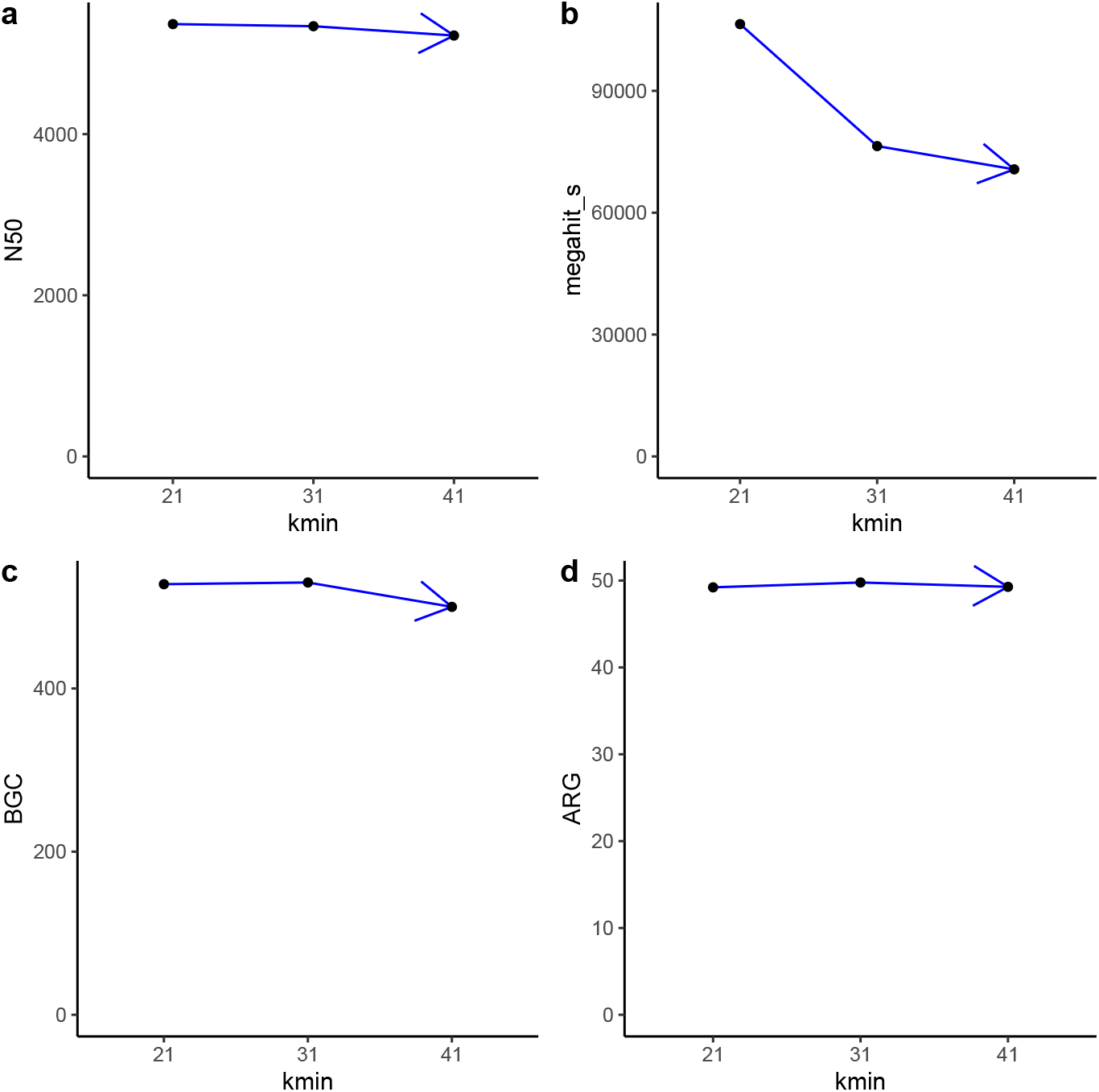
The impact of various settings for the *k-min* parameter on various metrics using the cave dataset. Each panel contains values for the *k-min* parameter on x-axis and the mean of a particular variable of interest grouped by the values of the *k-min* parameter on y-axis. **a** Comparison between the mean of N50 lengths by different *k-min* settings (left-top corner). **b** Comparison between the mean of assembly runtimes by different *k-min* settings (right-top corner). **c** Comparison between the mean of BGC counts by different *k-min* settings (left-bottom corner). **d** Comparison between the mean of ARG counts by different *k-min* settings (right-bottom corner).

**Figure 5.**
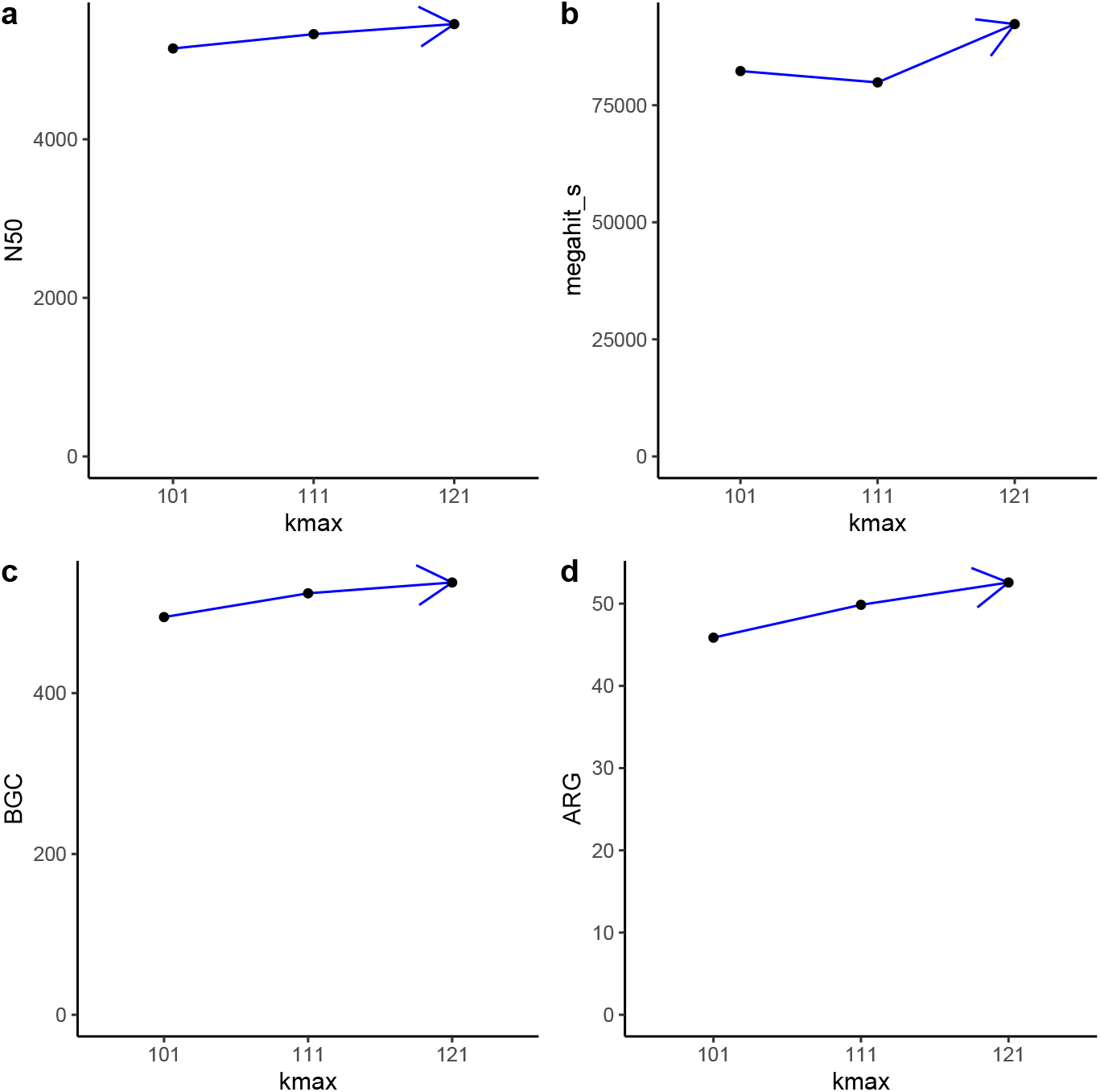
The impact of various settings for the *k-max* parameter on various metrics using the cave dataset. Each panel contains values for the *k-max* parameter on x-axis and the mean of a particular variable of interest grouped by the values of the *k-max* parameter on y-axis. **a** Comparison between the mean of N50 lengths by different *k-max* settings (left-top corner). **b** Comparison between the mean of assembly runtimes by different *k-max* settings (right-top corner). **c** Comparison between the mean of BGC counts by different *k-max* settings (left-bottom corner). **d** Comparison between the mean of ARG counts by different *k-max* settings (right-bottom corner).

### Results on wastewater metagenome samples

Similar to the previous experiment on the cave dataset, the same steps were performed using wastewater metagenome data. Figure 6 depicts Spearman’s rank correlations between the different variables of interest. Similar to Figure 1, a pair of variables (*total bp* and *contigs*) with a strong positive correlation (*r*_*S*_ ≈ 0.91) was encountered in Figure 6, but with a mild correlation to the BGC count in the case of both variables. However, fewer pairs of variables with corresponding rank correlation coefficients *>* 0.70 and pairs with negative correlations were detected.

**Figure 6.**
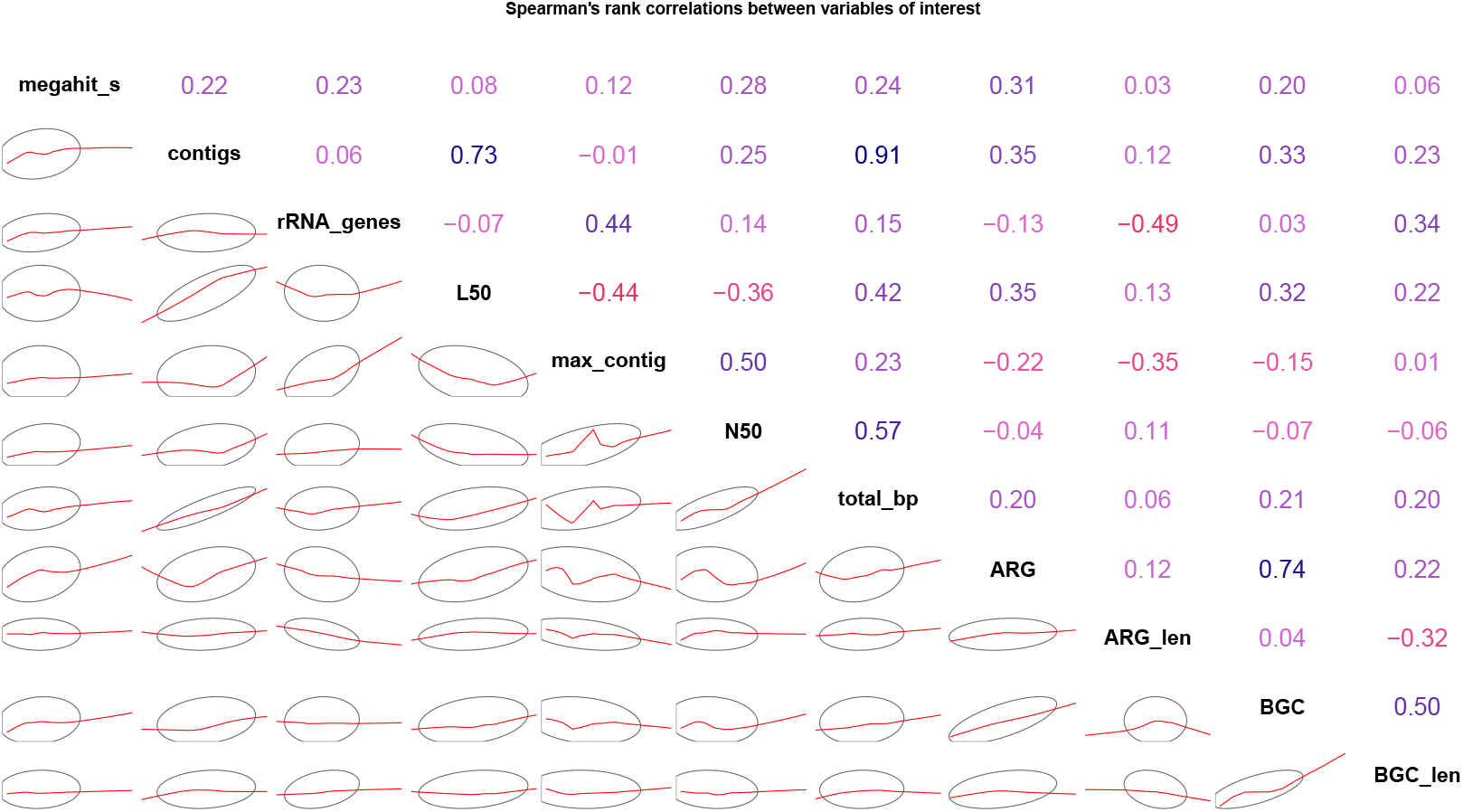
Spearman’s rank correlations between variables using the wastewater dataset. Variables are on the main diagonal; variable correlation coefficients for the whole dataset are on the upper-right panel; confidence ellipses and smoothed red lines, based on the corresponding correlation coefficient, indicating the overall 2D shape for each pair of two variables are shown on the lower-left panel. Blue color indicates positive correlation and red color indicates negative correlation, whereas the color intensity signifies the strength of the correlation.

After Friedman’s test (results shown in Table 5), the variables *ARG_len* and *BGC* were removed from further analysis because their corresponding p-value did not pass the significance threshold.

**Table 5.** P-values for assembly quality metrics computed using Friedman’s test for the wastewater dataset. The values were rounded to 3 decimals.

| Variable | p_value_Friedman |
| --- | --- |
| megahit_s | <0.001 |
| contigs | <0.001 |
| rRNA_genes | <0.001 |
| L50 | <0.001 |
| max_contig | <0.001 |
| N50 | <0.001 |
| total_bp | <0.001 |
| ARG | <0.001 |
| ARG_len | 0.429 |
| BGC | 0.164 |
| BGC_len | 0.003 |

The results of Conover’s all-pairs comparisons test for unreplicated blocked data are summarized in Fig. 7 in the form of a heatmap. For identifying significant differences easier, the p-values for pairwise scenario comparisons that were greater than 0.05 have been omitted from the figure. The p-values for pairwise comparisons that were less than or equal to 0.05 are marked with asterisks. The corresponding values of Conover’s significant statistics for all nine variables can be found in Table S8. Among the nine variables that were analyzed, *megahit_s* and *total_bp* showed significant differences in all 39 pairs of scenarios. Similarly, *contigs* showed significant pairwise differences in 30 of the 39 scenario pairs, while *N50* showed significant pairwise differences in 25 of the 39 scenario pairs. The *L50* and *rRNA_genes* variables had significantly different rank sum values in 12 and 7 of the 39 scenario pairs, respectively. The variable *ARG* showed significant pairwise differences in 16 of the 39 scenario pairs. However, *max_contig* and *BGC_len* did not show any significant difference in the pairs of scenarios involved.

**Figure 7.**
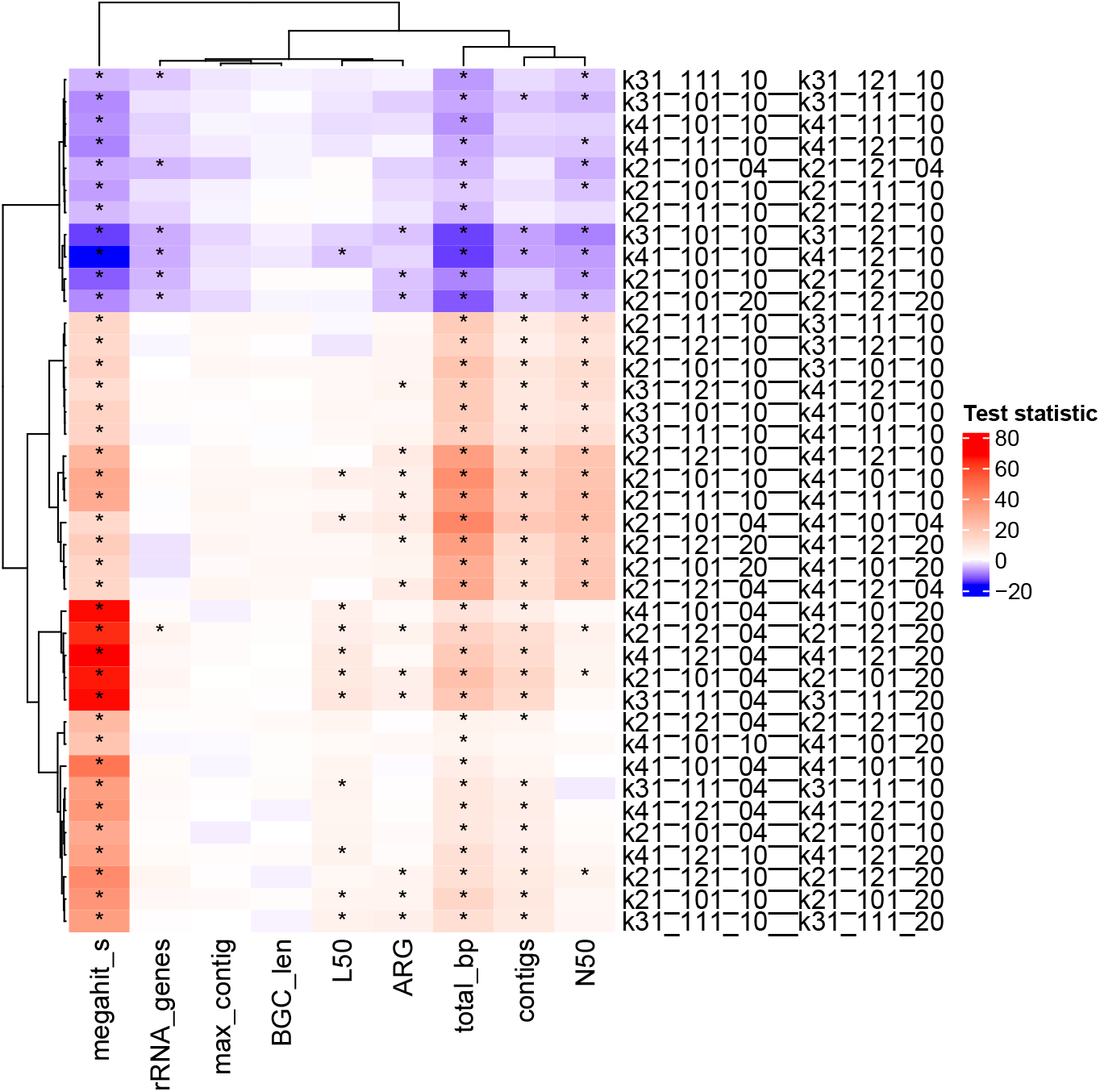
Heatmap showing test statistics from Conover’s all-pairs comparisons test for unreplicated blocked data, assessing differences across nine variables and thirty-nine pairwise scenario comparisons for the wastewater dataset. Red shades indicate that the first scenario in a pair had a higher rank sum (positive test statistic), while blue shades indicate the opposite (negative test statistic). Comparisons with p-values ≤0.05 are marked with asterisks. Rows (scenario pairs) and columns (variables) are hierarchically clustered using Euclidean distance to highlight patterns in statistical differences.

Regarding the hierarchical clustering of the pairwise scenario comparisons given as rows in Fig. 7, three main clusters were noticeable. These clusters of scenario pairs are presented in Table 6, where each column contains a list of scenario pairs that belong to a specific cluster. Specifically, cluster 1 contains 11 pairwise scenarios with varying *k-max*, cluster 2 consists of 13 pairwise scenarios with different values of *k-min*, and cluster 3 is composed of 15 pairwise scenarios with different *k-step*.

**Table 6.**
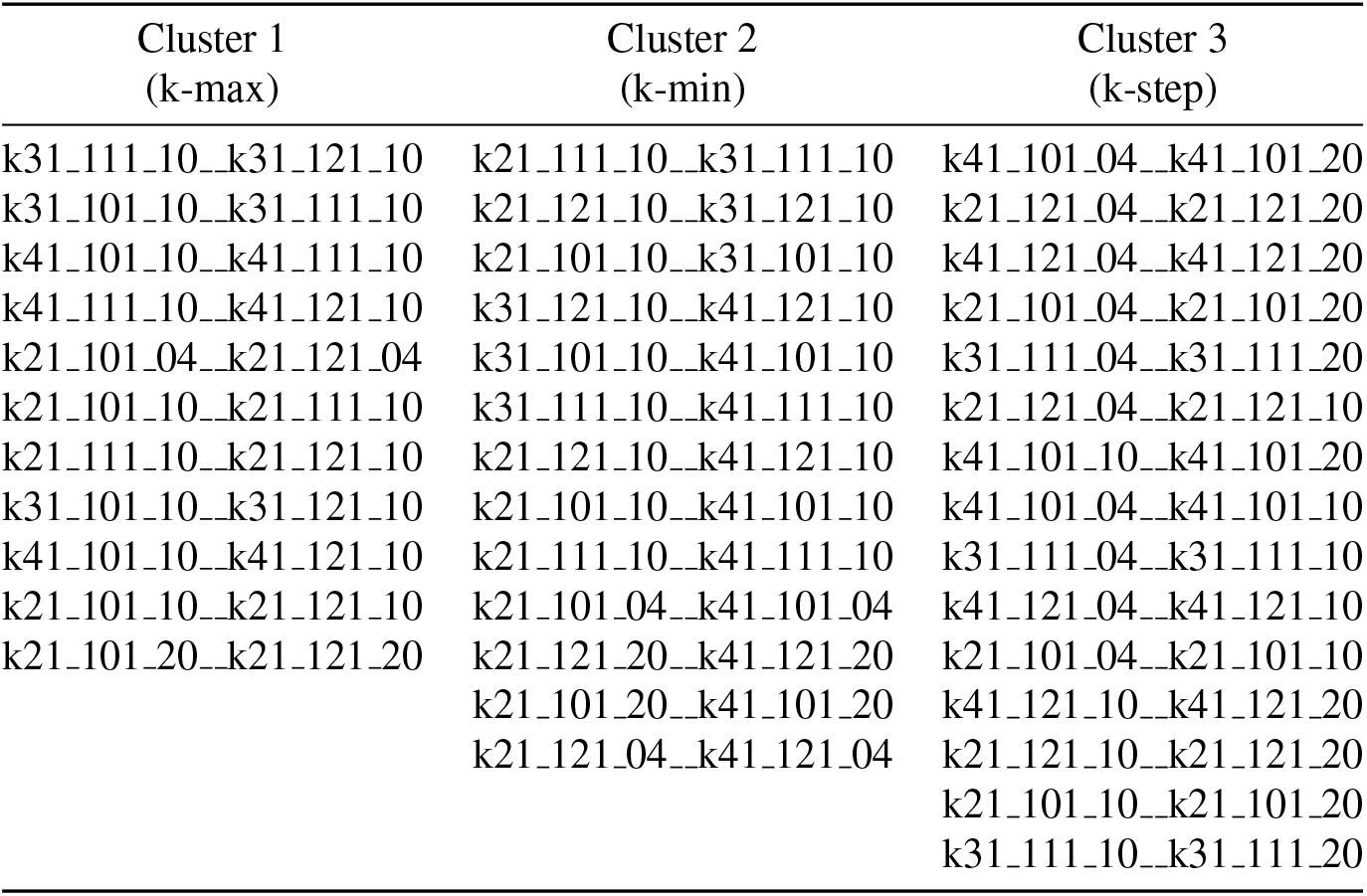
List of scenario pairs clustered using Euclidean distance for the wastewater dataset. Each column contains scenario pairs that belong to a specific cluster and only one k-mer parameter varies within each pair.

An additional grouping of the variables of interest (columns of Fig. 7) similar to the pairwise scenario clustering was provided and three main clusters were detected as follows. The first cluster was composed of one variable (i.e., *megahit_s*). The second cluster consisted of the variables *rRNA genes, max contig, BGC len, L50* and *ARG*. The third cluster contained the variables *total_bp, contigs* and *N50*.

The impact of different settings for the *k-step* parameter (values 4, 10, 20) on several quality metrics is presented in Fig. 8. Within this figure, each panel shows the *k-step* trade-off regarding a specific variable of interest, namely (a) N50 lengths, (b) assembly runtimes in seconds for MEGAHIT and (c) ARG counts, based on their mean value grouped by *k-step*. Similarly, the impact of different settings for the *k-min* parameter (values 21, 31, 41) on the same quality metrics was shown in Fig. 9. Further, the impact of different settings for the *k-max* parameter (values 101, 111, 121) on each of the three quality metrics (i.e., *N50, megahit_s* and *ARG*) is presented in Fig. 10. Compared to the previous analysis on cave data, the *BGC* variable was not included in these figures because Friedman’s test did not find any significant differences for this variable.

**Figure 8.**
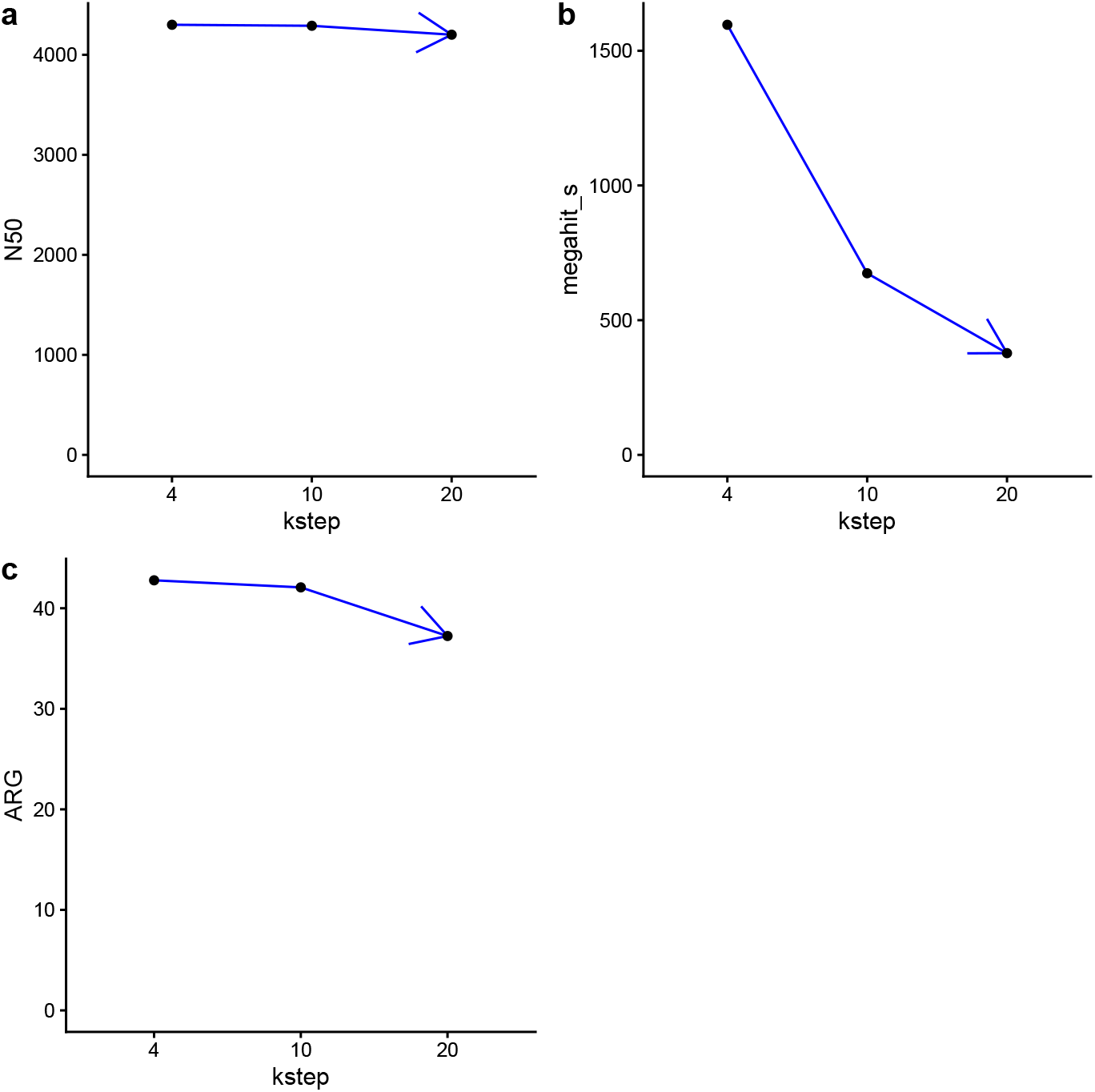
The *k-step* trade-off using the wastewater dataset where the x-axis contains values for the *k-step* parameter (i.e., 4, 10, 20). Values from the y-axis from each panel correspond to the mean of a particular variable of interest grouped by the values of the *k-step* parameter. **a** Comparison between the mean of N50 lengths by different *k-step* settings (left-top corner). **b** Comparison between the mean of assembly runtimes in seconds using MEGAHIT by different *k-step* settings (right-top corner). **c** Comparison between the mean of ARG counts by different *k-step* settings (left-bottom corner).

**Figure 9.**
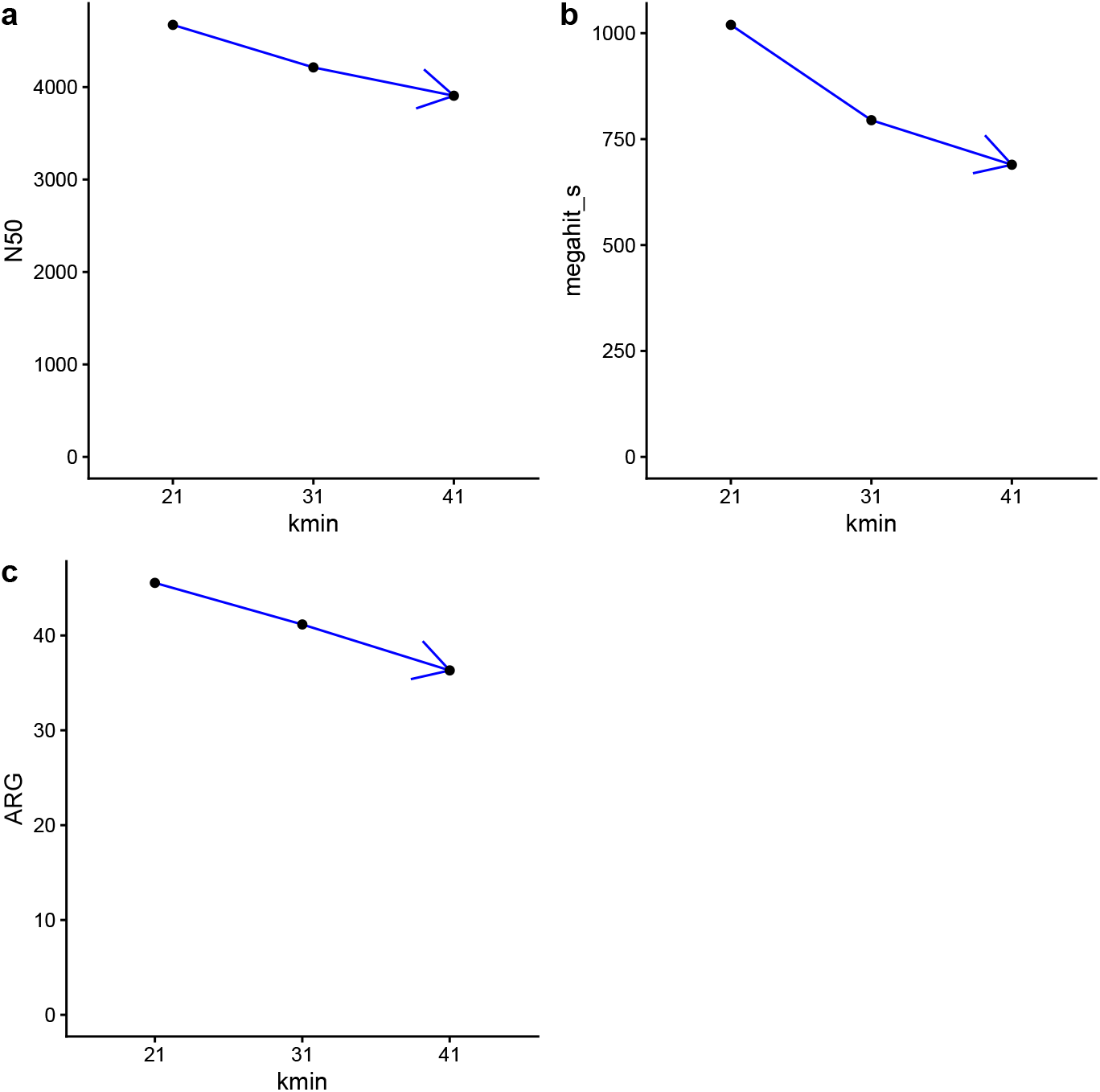
The impact of various settings for the *k-min* parameter on various metrics using the wastewater dataset. Each panel contains values for the *k-min* parameter on x-axis and the mean of a particular variable of interest grouped by the values of the *k-min* parameter on y-axis. **a** Comparison between the mean of N50 lengths by different *k-min* settings (left-top corner). **b** Comparison between the mean of assembly runtimes by different *k-min* settings (right-top corner). *c* Comparison between the mean of ARG counts by different *k-min* settings (left-bottom corner).

**Figure 10.**
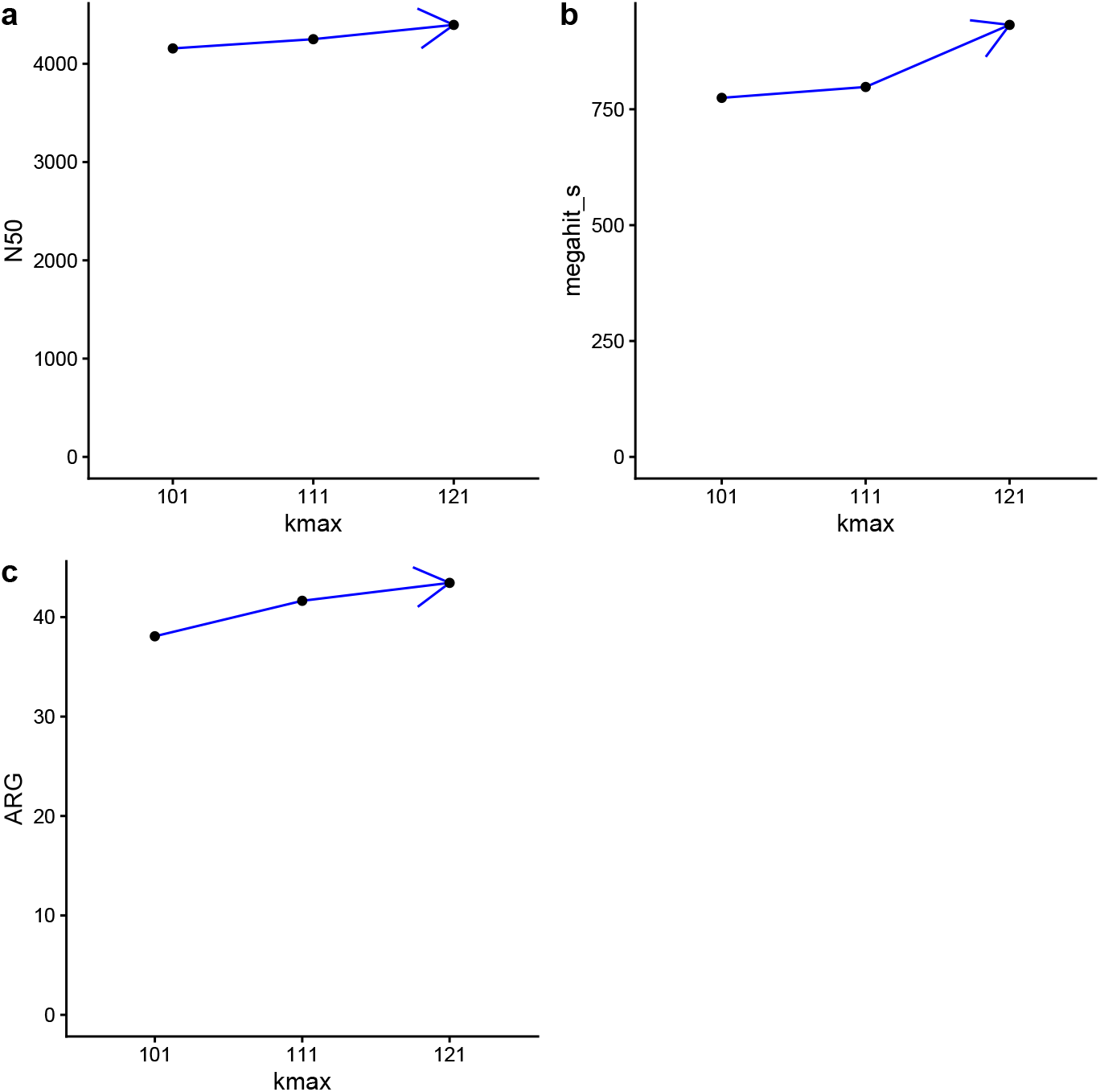
The impact of various settings for the *k-max* parameter on various metrics using the wastewater dataset. Each panel contains values for the *k-max* parameter on x-axis and the mean of a particular variable of interest grouped by the values of the *k-max* parameter on y-axis. **a** Comparison between the mean of N50 lengths by different *k-max* settings (left-top corner). **b** Comparison between the mean of assembly runtimes by different *k-max* settings (right-top corner). **c** Comparison between the mean of ARG counts by different *k-max* settings (left-bottom corner).

## DISCUSSION

In this study, we investigated the role of the k-mer size ranges in MEGAHIT on metagenomic assembly in the context of microbial community-wide identification of ARGs and BGCs from cave and wastewater microbiomes. We observed that different k-mer settings were associated with differences in the corresponding values for variables of interest from one scenario to another. Only pairs of scenarios with a single difference at the level of *k-min, k-max* or *k-step* were considered for further analysis.

### Analysis of cave dataset

For the cave metagenome samples specifically, we observed that the pairs of scenarios that were compared revealed different patterns in terms of test statistic values and p-values computed via Conover’s all-pairs comparisons test for unreplicated blocked data (see Fig. 2 and Table S7). Scenarios with smaller *k-step* had increased runtimes compared to scenarios with larger values for this parameter (and a smaller corresponding number of k-mer sizes in their range).

This trade-off between different values for the *k-step* parameter is also observable in Fig. 3, where the selection of the *k-step* value had an impact on various assembly metrics such as N50 lengths, assembly runtime in seconds, BGC counts or ARG counts. More specifically, as *k-step* increased, the runtime also decreased significantly, but this efficiency gain implied a trade-off of reduced BGC and ARG recovery, while N50 showed a similar but less impactful change. In contrast, when *k-max* increased, the values for BGC counts, ARG counts and *N50* increased, with a trade-off of longer assembly runtimes (Fig. 5).

Similarly, the effect of different settings for the *k-min* parameter on these four metrics is shown in Fig. 4. Notably, for fourteen out of fifteen pairs of scenarios (with the exception of k41_101_10_k41_101_20) with varying *k-step*, the scenarios with smaller *k-step* also showed an increased number of assembled contigs and identified BGCs, while in terms of *N50*, which is a way of expressing the level of fragmentation of an assembly, only three pairs showed a significant difference.

For pairs of scenarios with different values for either *k-min* or *k-max*, the results were mixed. Based on the hierarchical clustering of pairwise scenarios in Fig. 2, most pairs belonging to the first cluster had significant differences detected in the second scenario from pairwise comparison, where only the value of the parameter *k-max* varied. Scenarios with larger *k-max* but the same *k-min* and *k-step* also had an increased assembly runtime, except for five pairs (i.e., k21_111_10_k21_121_10, k31_111_10_k31_121_10, k21_101_10_k21_111_10, k41_111_10_k41_121_10 and k21_101_04_k21_121_04), which did not show significant differences in terms of runtime (*megahit_s*). Furthermore, when only *k-max* varied for eleven pairs of scenarios (listed in Table S4), the larger value of this parameter was associated with a significantly increased N50 for all pairs except k21_111_10_k21_121_10 and the number of assembled contigs for all pairs except k31_111_10_k31_121_10, in addition to an increased assembly runtime for six of these scenario pairs. This suggests that it may be possible, in principle, to further optimize these assembly quality metrics by further increasing the *k-max*. However, in particular, the current version of MEGAHIT imposes an upper bound of 127 for the *k-max* parameter.

Within the second cluster of scenario pairs characterized by different values for *k-step*, four pairs of scenarios showed significant differences for the first scenario in terms of *rRNA genes, ARG, megahit_s, contigs, L50, total_bp* and *BGC* (i.e., k21_101_04_k21_101_20, k21_121_04_k21_121_20, k31_111_04_k31_111_20 and k41_121_04_k41_121_20). An exception (i.e., k41 101 04 k41 101 20) did not show significant differences in terms of the number of identified ARGs, but showed significant differences in terms of *rRNA genes, megahit_s, contigs, L50, total_bp* and *BGC*. For the other variables of interest, the results were mixed for this cluster.

Conversely, when only *k-min* varied for thirteen pairs of scenarios (listed in Table S3), an observed difference was in terms of assembly runtime for the first scenario for all pairs, suggesting that starting the k-mer range from a lower value did not bring a significant change in terms of the variables of interest.

Similar to the scenario pairs with different values for *k-step*, these scenario pairs were split between the third and the fourth cluster, respectively.

Specifically, the third cluster of pairs of scenarios was characterized by different values of *k-min* for the scenarios in each pair (Table 4). All pairs showed an increased assembly runtime associated with the first scenario from pairwise comparisons. However, the variables *rRNA genes* and *total_bp* had significant differences for the pair of scenarios k21_121_20_k41_121_20, namely for the second and first scenarios, respectively. Furthermore, the pair of scenarios k21_101_20_k41_101_20 also showed significant differences in terms of *BGC len* for the first scenario of the pair. However, none of these pairs indicated significant differences in terms of the number of ARGs and BGCs (Fig. 2).

Compared to the previous clusters of pairwise scenario comparisons, the fourth cluster consisted of pairs with different *k-step* and pairs with different *k-min*. Most of these scenario pairs showed significant differences in terms of *megahit_s, contigs, L50, total_bp* and *BGC*, while larger values for these variables were associated with reduced values for *k-min* or *k-step*. These scenario pairs can be grouped into 4 pairs with different *k-min* and 9 pairs with different *k-step*, based on the k-mer parameter that varies. However, there were no significant differences for the variable *L50* using a reduced *k-min* in the case of four pairs of scenarios (i.e., k21_101_10_k41_101_10, k21_111_10_k41_111_10, k21_101_04_k41_101_04 and k21_121_04_k41_121_04), while one pair (i.e., k41_101_10_k41_101_20) showed significant differences in terms of *megahit_s, contigs* and *L50* within the first scenario with a reduced *k-step*. Meanwhile, the results for the other variables of interest (*N50, max_contig, BGC_len, rRNA_genes* and *ARG*) among these scenario pairs were inconsistent.

Additionally, a summary of the significant effects on assembly and gene discovery metrics by applying different settings for k-mer parameters, grouped by clusters, was provided in Table S9. When *k-max* increased, significant differences were detected for scenarios with a larger value for this parameter in terms of assembly metrics (e.g., *N50, megahit_s, contigs*) and gene discovery metrics (*BGC* - 10 pairs and *ARG* - 5 pairs). When *k-step* increased, the corresponding scenarios showed significant differences in terms of *N50* (3 pairs), while scenarios with a smaller *k-step* value yielded more ARG counts (8 pairs), longer assembly runtimes (all 15 pairs), more contigs within assemblies (all 15 pairs) and more BGCs detected (14 pairs). When *k-min* increased, scenarios with a larger *k-min* value showed an increased value for variables *max_contig* (1 pair), *rRNA_genes* (1 pair), while scenarios with a smaller *k-min* value showed an increased value for variables *N50* (7 pairs), *ARG* (1 pair), *megahit_s* (all 13 pairs), *contigs* (8 pairs) and *BGC* (8 pairs).

Some scenarios that we envisaged, namely those with a *k-step* of 20 (the largest *k-step* value considered here), are comparable to the *reduced set of k-mers* from the work of Qayyum et al. (2025). In their study, the authors concluded that the reduced set of k-mers achieved similar results (in terms of various assembly quality metrics) to the results of scenarios with larger sets of k-mers (with reduced *k-step*), while significantly reducing the runtime of MEGAHIT. The previously mentioned k-mer scenarios were involved in several pairwise comparisons in our study (see Table S6). Among these, all pairs of scenarios showed significant differences in terms of the number of contigs from the resulting assemblies. In addition, all pairs of scenarios (except k41_101_10_k41_101_20) also showed a significantly reduced value for *BGC* in the scenario with a larger *k-step*.

Furthermore, an increased *k-step* for scenarios also led to a significant reduced value for *ARG* counts in the case of eight out of fifteen scenario pairs with varying *k-step* values (Fig. 2). Besides these, among the fifteen pairwise comparisons with varying *k-step* value, there were pairs that showed significant differences in terms of at least one of the other variables of interest such as *N50* (3 pairs), *max_contig* (2 pairs), *BGC_len* (2 pairs), *rRNA_genes* (10 pairs), *L50* (all 15 pairs) or *total_bp* (14 pairs) (Fig. 2). This suggests that when ARGs and BGCs are also of interest, in addition to assembly quality, more care should be taken to identify suitable k-mer ranges for MEGAHIT, in order to optimize the identification of these elements.

### Analysis of wastewater dataset

Compared to the previous experiment using cave data (Fig. 2), we observed a similar trend regarding the effect of the k-mer parameter settings on the nine variables of interest (Fig. 7). Additionally, the information about the significant effect of the k-mer parameter settings from Fig. 7 is presented in Table S10. Here, the variable *BGC* was removed from the Conover’s test as a consequence of its lack of statistically significant results after Friedman’s test, as previously mentioned. These scenario pair comparisons uncovered different patterns in terms of test statistic values as well as p-values computed via Conover’s all-pairs comparisons test (Fig. 7). Specifically, scenarios with smaller *k-step* also had an increased runtime compared to the scenarios with larger values for this parameter. Beside this, increasing the *k-step* parameter also led to fewer discoverable ARGs (6 scenario pairs), a smaller number of contigs (13 scenario pairs), a smaller value for *total_bp* (15 scenario pairs), fewer rRNA_genes (1 scenario pair), a smaller value for *L50* (9 scenario pairs) and a smaller value for *N50* (5 scenario pairs). However, variables *max_contig* and *BGC_len* were not significantly affected by increasing the *k-step* parameter. All these pairwise scenario comparisons were mentioned in the third column of Table 6 and in Table S5, respectively. This trade-off between different values for the *k-step* parameter in terms of *N50, megahit_s* and *ARG* is also observable in Fig. 8.

When comparing scenarios with different *k-max*, scenarios with larger *k-max* showed an increased assembly runtime (11 scenario pairs), more ARG counts (3 scenario pairs), more rRNA_genes (6 scenario pairs), a larger value for *L50* (1 pair), a larger value for *total_bp* (11 scenario pairs), a larger number of contigs (4 scenario pairs), a larger value for *N50* (9 scenario pairs). Additionally, increasing *k-max* did not show any significant differences for *max_contig* and *BGC_len*. All scenario pairs with different *k-max* were enumerated in the first column of Table 6 and in Table S4. The impact of the selection for *k-max* on *N50*, assembly runtime and ARG counts is shown in Fig. 10.

In terms of varying *k-min*, scenarios with smaller *k-min* showed an increased assembly runtime (13 scenario pairs), more ARGs (7 scenario pairs), a larger value for *L50* (2 scenario pairs), a larger value for *total_bp* (13 scenario pairs), a larger number of contigs (13 scenario pairs), a larger value for *N50* (13 scenario pairs). However, no significant differences were detected for *rRNA_genes, max_contig* and *BGC_len* when increasing the *k-min* parameter. All scenario pairs with different *k-min* were mentioned in the second column of Table 6 and in Table S3. The effect of different settings for the *k-min* parameter on three quality metrics (*N50, megahit_s, ARG*) is presented in Fig. 9.

Among these 39 scenario pairs, there were 10 pairs with different *k-step*, containing one scenario with the largest value for this parameter (i.e., 20) (Table S6). These scenario pairs were used to test whether a reduced set of k-mer sizes proposed by Qayyum et al. (2025) could have an impact on the variables, including the number of ARG and BGC counts. Within these pairs, the scenario with an increased *k-step* showed a reduced assembly runtime (i.e., *megahit_s*), but also a reduced number of contigs (*contigs*) for all pairs except k41_101_10_k41_101_20. Here, the scenario pair k21_121_04_k21_121_20 had a smaller value for *rRNA_genes* for the scenario with an increased *k-step*.

Furthermore, increasing the *k-step* parameter had no impact on variables *max_contig* and *BGC_len*. Regarding the degree of assembly contiguity represented by *N50*, using the reduced set of k-mer sizes was associated with higher N50 values for only 3 scenario pairs (i.e., k21_121_04_k21_121_20, k21_101_04_k21_101_20 and k21_121_10_k21_121_20). In terms of *L50*, the scenario with a smaller *k-step* value had a larger corresponding value for this variable in the case of 8 scenario pairs (i.e., k41_101_04_k41_101_20, k21_121_04_k21_121_20, k41_121_04_k41_121_20, k21_101_04_k21_101_20, k31_111_04_k31_111_20, k41_121_10_k41_121_20, k21_101_10_k21_101_20 and k31_111_10_k31_111_20). In addition to these, a significant effect of the *k-step* parameter on variable *total_bp* was detected within the scenario with a smaller value for this parameter in all 10 scenario pairs. In terms of ARG counts, increasing the *k-step* parameter led to the detection of fewer ARGs, while the scenario with a smaller *k-step* had a significant impact on variable *ARG* in 6 scenario pairs (i.e., k21 121 04 k21 121 20, k21 101 04 k21 101 20, k31 111 04 k31 111 20, k21 121 10 k21 121 20, k21 101 10 k21 101 20 and k31 111 10 k31 111 20). The other scenario pairs with varying *k-step*, which included a scenario with *k-step*=20, did not show significant differences in terms of ARG counts.

### Comparative analysis between caves and wastewater dataset results

In this study, two different environments, namely karstic caves and a wastewater system, were studied to identify functional genes (ARGs and BGCs). In terms of samples, 17 sediment samples collected from 4 sites (Cloşani Cave, Ferice Cave, Bears’ Cave and Movile Cave from Romania) were used for the first analysis, while 10 wastewater samples from a campus wastewater system were used in the second analysis. The same experimental setup was applied for both datasets in terms of k-mer length parameter settings (for assembly scenarios), bioinformatics tools, pipelines and evaluation metrics.

The two datasets showed the same main trade-off: coarser k-mer ranges had reduced runtime, whereas finer-grained settings preserved more biological signal. In cave samples, this pattern was strongest for *BGC* recovery and was also visible for *ARG* counts. In wastewater samples, the clearest gene-level effect was on *ARG* recovery, while *BGC* did not remain significant after the overall Friedman test. Across both datasets, increasing k-max generally increased *N50* and runtime. By contrast, the effect of *k-min* was less uniform across datasets and should be interpreted more cautiously than the *k-step* and *k-max* effects.

Another difference between these two datasets is given by the way these 39 scenario pairs were classified based on Euclidean distance using the Conover’s test. Specifically, the scenario pairs were grouped into 4 clusters for the first experiment (cave data) and 3 clusters for the second experiment (wastewater data). The quality metrics were also classified based on Euclidean distance (2 clusters for the first experiment and 3 clusters for the second experiment).

Possible limitations of these experiments include the relatively limited number of samples per dataset, which may reduce precision. In addition, because total assembly length and contig count were strongly correlated with BGC count in the cave dataset, higher BGC recovery may be interpreted as an increased predicted signal after assembly, not as an independent estimate of true environmental BGC abundance. Using two distinct environments strengthens the evidence that the runtime/ARG trade-off was not restricted to a single dataset type. However, the BGC result should be interpreted more cautiously: it was clear in the cave dataset, but BGC did not remain significant after Friedman’s test in the wastewater dataset, possibly related to the lower sequencing depth of the wastewater subset, although differences in environment, community composition and sample selection may also have contributed. This caveat is consistent with the reported sequencing depth in the supplementary tables: the wastewater subset had about 0.67 Gb per sample on average, whereas the cave subset had about 60.6 Gb per sample on average, as shown in Tables S1 and S2. Additional datasets from other environments will be needed before extending these findings to higher-complexity metagenomes.

Another limitation of this kind of study, in general, is that dedicated access to computing resources needs to be ensured in order to provide a fair comparison of runtimes across different scenarios. However, this limitation is common for this type of study, when runtime is being taken into account.

## CONCLUSIONS

This study revealed that the selection of k-mer parameters in MEGAHIT is a critical decision that introduces a trade-off between computational cost and functional gene discovery. Additionally, the functional gene discovery also depends on the input data (sequenced reads) for the assembly tool. Based on our findings, we offer the following recommendations: (1) When aiming to maximize the discovery of ARGs and BGCs, researchers should consider using fine-grained k-mer sets (i.e., *k-step* ≤ 4 with *k-max* close to its upper limit) in similar discovery-oriented analyses, accepting longer assembly times as the cost of a more complete assembly and maximizing the recovery of functional elements. (2) If computational resources are limited and ARG or BGC discovery is not the primary aim, coarser k-mer ranges can be used, but with the caveat that some ARGs and BGCs may be missed.

Future work would be required in order to replicate these results on a variety of metagenomic datasets from diverse environments, including a wider range of sample types. In addition, future studies could envisage the implementation of optimization algorithms in order to maximize or minimize the values of the desired quality metrics relative to the 3 parameters that control the k-mer range in MEGAHIT.

## AUTHOR CONTRIBUTIONS

- Alina Cărunta conceived and designed the experiments, performed the experiments, analyzed the data, prepared figures and/or tables, authored and reviewed the drafts of the article, and approved the final draft.
- Horia Leonard Banciu provided the raw sequenced reads from the cave samples, conceived and designed the experiments, authored and reviewed the drafts of the article, and approved the final draft.
- Alexandru Eugeniu Mizeranschi conceived and designed the experiments, analyzed the data, prepared figures and/or tables, authored and reviewed the drafts of the article, and approved the final draft.

